# Cellular Density Homeostasis Is Governed by an Anisotropic Attractor and Predictive Feedback Control

**DOI:** 10.64898/2026.09.28.755031

**Authors:** Victor Alcolea-Rodriguez, Anoushka Gupta, Michael Ka Ho Lee, Dario Polli, Renzo Vanna, Ishan Barman

## Abstract

Cell density is a fundamental biophysical property of living cells, yet the dynamical architecture underlying density homeostasis remains unknown. Here, we reconstruct the geometry, dynamics, and control structure governing cellular density regulation from single-cell mass–volume measurements acquired during recovery from osmotic perturbation. We identify a persistent anisotropic attractor in mass–volume state space that permits coordinated density-preserving fluctuations while strongly suppressing density-changing deviations. Optimal transport analysis reveals directed recovery trajectories confined to this attractor, enabling inference of an effective density-restoring control law. Density deviations are associated with coordinated changes in biomass accumulation and volume expansion that together generate an effective density-restoring control law. Moreover, density error contains significant predictive information regarding future corrective behavior, revealing an information-rich component of cellular regulation. Together, these findings establish cellular density homeostasis as an emergent dynamical control process and demonstrate how attractor reconstruction, optimal transport, and information theory can be combined to uncover the organizing principles of biological homeostasis.

## Introduction

The ability to maintain stable internal states despite continuous perturbations is a defining characteristic of living systems.^1^ From metabolic regulation and osmotic balance to developmental patterning and tissue organization, biological function depends on homeostatic mechanisms that preserve critical physiological variables while remaining sufficiently adaptable to changing environments.^2,3^ Although homeostasis has traditionally been studied through the lens of molecular pathways and regulatory networks, increasing evidence suggests that many biological systems can be understood as dynamical systems governed by feedback control.^4–6^ Within this framework, stable physiological states emerge as attractors in an underlying state space, while robustness arises from coordinated restoring dynamics that counteract perturbations^7,8^. Such concepts have successfully explained diverse phenomena ranging from bacterial chemotaxis^9^ and cell-size regulation^10^ to neural activity^11^ and developmental fate decisions^12^. Yet despite the central importance of homeostasis in biology, the dynamical architectures underlying many homeostatic processes remain poorly characterized.

Cellular density represents an especially intriguing example of biological homeostasis. Defined as the ratio of dry mass to volume, density integrates the cumulative effects of biosynthesis, degradation, metabolism, growth, and osmotic regulation.^13^ Because density influences macromolecular crowding, intracellular transport, reaction kinetics, and mechanical properties, its maintenance is essential for cellular function.^14,15^ Remarkably, many cell types preserve density within a relatively narrow range despite substantial fluctuations in mass accumulation and water content.^16^ This observation suggests the existence of active regulatory mechanisms capable of detecting density deviations and generating corrective responses.^17^ However, while density homeostasis has become increasingly recognized as a fundamental cellular property,^13^ the principles governing its regulation remain incompletely understood.

Recent advances in quantitative phase imaging (QPI) have transformed the study of cellular growth and biophysics by enabling label-free measurements of single-cell dry mass and volume with high precision.^14,18,19^ These approaches have revealed unexpected complexity in the regulation of cell size, growth, and density, highlighting the importance of single-cell heterogeneity and dynamical behavior.^20,21^ Using QPI, we recently demonstrated that mammalian cells subjected to osmotic perturbation actively restore their pre-perturbation density through coordinated but distinct mass and volume recovery processes.^14^ These findings established density homeostasis as a robust and regulated cellular phenomenon and raised a broader question: what dynamical principles enable cells to maintain and restore density following perturbation?

Addressing this question requires moving beyond measurements of average behavior and toward reconstruction of the state-space geometry underlying density regulation. If density is actively controlled, then cells should not occupy mass–volume space randomly. Instead, populations should organize around preferred regions of state space, deviations from these regions should generate restoring dynamics, and recovery trajectories should reveal signatures of feedback control.^10^ Such ideas have proven powerful in developmental biology,^12^ neuroscience,^11^ and systems biology,^22^ where attractor landscapes provide intuitive descriptions of robustness, stability, and state transitions. However, analogous dynamical descriptions of density homeostasis have remained largely unexplored.

A major challenge is that the trajectories governing homeostatic regulation are often difficult to observe directly. While long-term tracking of individual cells can provide important insights, homeostatic behavior frequently emerges at the population level and may involve heterogeneous trajectories that are difficult to reconstruct from single-cell longitudinal observations alone. Recent developments in optimal transport theory provide a powerful framework for addressing this challenge.^23^ Rather than requiring continuous tracking of individual cells, optimal transport reconstructs a minimum-cost coupling of probability mass between experimentally observed population states, enabling inference of effective vector fields and dynamical trajectories from time-resolved snapshot measurements.^24,25^ Originally developed in mathematics and economics and more recently applied to developmental and single-cell genomic systems,^26^ optimal transport offers a natural framework for reconstructing the dynamics of biological homeostasis from population-scale measurements.^27^

Beyond geometry and dynamics, homeostatic regulation can also be viewed as an information-processing problem. Effective controllers must not only detect deviations from a target state but also generate corrective responses whose magnitude and direction depend on the nature of the perturbation. From this perspective, the performance of a homeostatic system is reflected not only in its ability to suppress fluctuations but also in the predictive information contained within its regulatory variables. Information-theoretic approaches have provided important insights into sensory adaptation,^28^ signaling networks,^29^ and biological feedback control,^9^ yet they have rarely been applied to the problem of cellular density regulation.

Here, we revisit the single-cell mass–volume measurements underlying our previous study of density recovery through the lens of dynamical systems theory, optimal transport, and information theory. This perspective enables reconstruction of the state-space geometry, restoring dynamics, and feedback architecture underlying density homeostasis, revealing principles that were not accessible through conventional analyses of mass and volume recovery alone. We demonstrate that cell populations occupy a low-dimensional attractor embedded within mass–volume space and that recovery following osmotic perturbation is governed by highly structured restoring dynamics. Optimal transport analysis reveals a directed flow field that drives cells toward the attractor, enabling inference of an effective density-restoring control law. Finally, we show that corrective behavior becomes increasingly likely as cells move farther from the attractor and that density error contains measurable predictive information about future corrective responses. Together, these findings establish cellular density homeostasis as an emergent dynamical control process governed by attractor geometry, restoring dynamics, and information-rich feedback regulation, and provide a general framework for reconstructing biological control systems from single-cell measurements. More broadly, this work illustrates how state-space reconstruction, optimal transport, and information theory can be integrated to infer the geometry, dynamics, and control architecture of biological homeostatic systems directly from experimental measurements.

## Results

### Cellular density homeostasis emerges from coordinated mass–volume recovery following osmotic perturbation

To investigate the dynamical organization of cellular density homeostasis, we analyzed single-cell dry mass and volume measurements of NIH-3T3 fibroblasts acquired at multiple timepoints following osmotic perturbation (0, 1, 4, 7, 24, and 48 h). These measurements provide a quantitative description of how cell populations move through mass–volume state space during recovery.

Across all timepoints, dry mass and volume remained strongly coordinated, with cells occupying a narrow region of state space organized around a characteristic linear relationship corresponding to a target cell mass density (CMD) of *ρ*₀ = 182.1 fg/μm³ (**Fig. 1A**). Despite substantial physiological heterogeneity and perturbation-induced changes in population composition, cells remained closely aligned with this CMD manifold.

**Figure 1.**
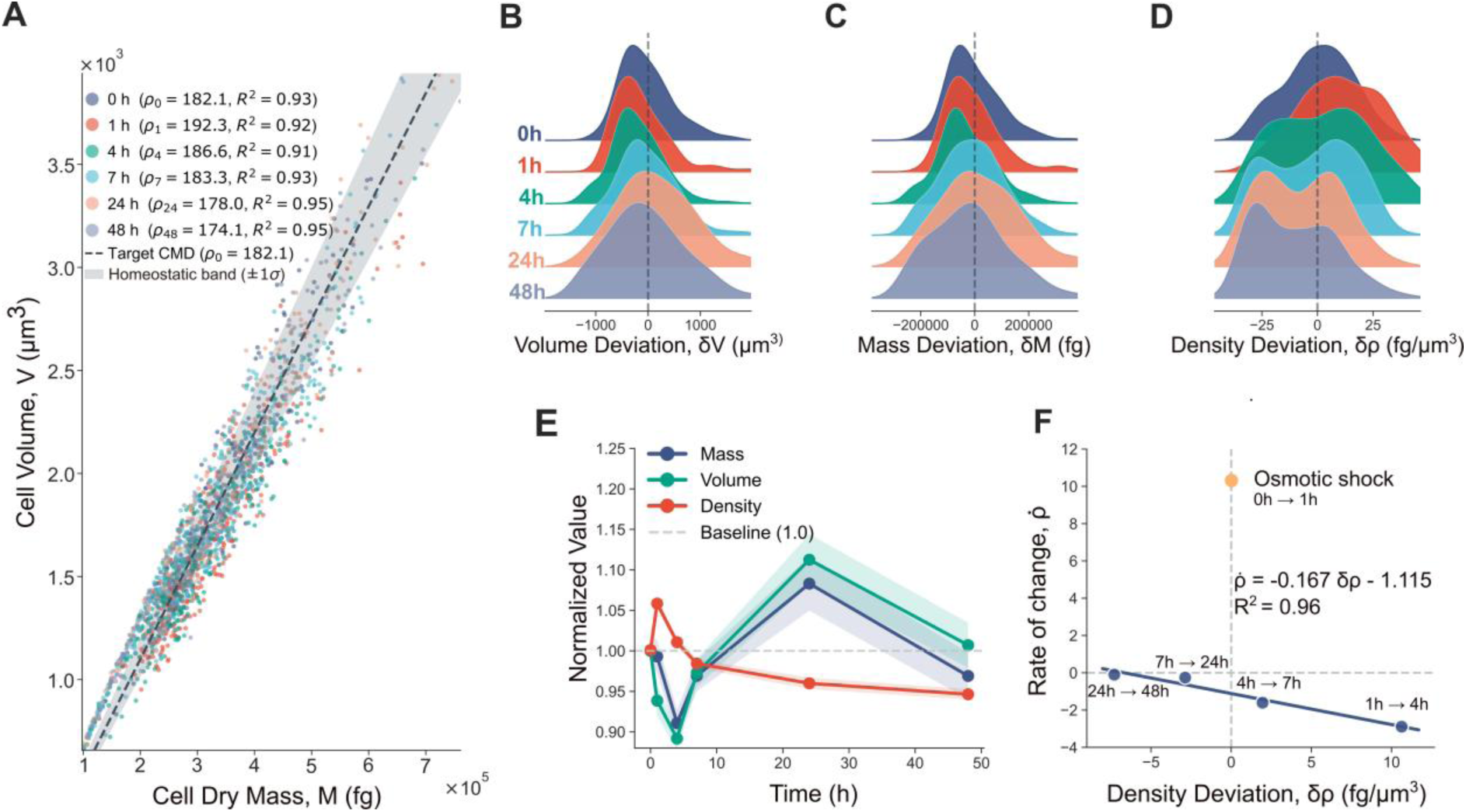
Cellular density homeostasis emerges from coupled mass-volume dynamics following osmotic shock. (A) Single-cell scaling of dry mass versus volume during a 48-h recovery. Cells remain tightly aligned with the target cell mass density (CMD; ρ₀ = 182.1 fg/μm³, dashed line) and within the ±1σ homeostatic band (shaded) across all timepoints (R² ≥ 0.91). (B-D) Temporal population distributions of deviations in (B) volume (δV), (C) mass (δM), and (D) density (δρ). While mass and volume exhibit broad transient shifts, density is constrained and progressively returns to the homeostatic setpoint (zero-deviation line). (E) Population-averaged trajectories of mass, volume, and density normalized to pre-perturbation values. Shaded regions indicate standard error mean (SEM). (F) State-space dynamics of density restoration. The initial osmotic shock (0-1 h, star) corresponds to the acute perturbation-induced density increase. During recovery (1-48 h), the rate of density change (ρ̇) exhibits a strong negative linear dependence on density deviation at the beginning of each interval (R² = 0.96).

The osmotic shock triggered coordinated deviations in both dry mass and volume (**Fig. 1B, 1C, 1E**). Following this transient state-space displacement, the population density distribution progressively relaxed toward the baseline state (**Fig. 1D**). This trajectory demonstrates that density homeostasis is not achieved by constraining mass or volume independently, but rather emerges from their coordinated co-regulation.

To rigorously assess whether density recovery operates via active feedback, we analyzed the relationship between the density deviation (δρ, as a result of *ρ_t_* - *ρ*₀) and its subsequent rate of change (ρ̇). The 0–1 h interval, corresponding to the acute osmotic perturbation, produced a marked increase in population-averaged density at an average rate of +10.2 (fg/μm³)/h and was therefore analyzed separately from the recovery phase (**Fig. 1F, yellow dot**). In contrast, the subsequent biological cell population exhibited a robust linear dependence of ρ̇ on δρ (R² = 0.96). High-density states generated strong negative rates of change, which progressively attenuated as cells approached the target density, thereby mediating a stable, restorative return to the homeostatic setpoint.

Together, these observations establish three key features of the system. First, cellular populations remain organized around a preferred density manifold with coordinated substantial fluctuations in mass and volume. Second, density deviations remain bounded and progressively recover following perturbation. Third, recovery exhibits a restoring tendency consistent with active homeostatic regulation. These findings suggest that density homeostasis may be governed by an underlying dynamical structure within mass–volume state space, motivating reconstruction of the attractor geometry underlying this behavior.

### Density homeostasis is associated with an anisotropic attractor in mass–volume state space

The persistent organization of cells around a common CMD manifold suggested that density homeostasis may be governed by an underlying empirical attractor in mass–volume state space. To investigate the geometry governing density homeostasis, we reconstructed empirical quasi-potential landscapes (*U*) directly from the observed population distributions in mass–volume space. Here, *U* does not represent a thermodynamic potential or imply conservative cellular dynamics; rather, it serves as a quantitative metric of state-space occupancy, where highly populated states correspond to low quasi-potential regions, whereas sparsely occupied states correspond to higher quasi-potential regions

The reconstructed landscape at baseline (t = 0 h) revealed a pronounced valley extending along the CMD manifold (**Fig. 2A and 2B**). Rather than converging to a single preferred mass or volume, cells occupied an elongated low-potential region spanning a broad range of physiological states. Following osmotic perturbation (t = 1 h), the population distribution shifted within state space, producing a transient displacement of the fitted density axis relative to the target CMD (**Fig. 2A**). Despite this perturbation, the overall landscape topology remained remarkably preserved, with the dominant low-potential valley continuing to align closely with the CMD manifold.

**Figure 2.**
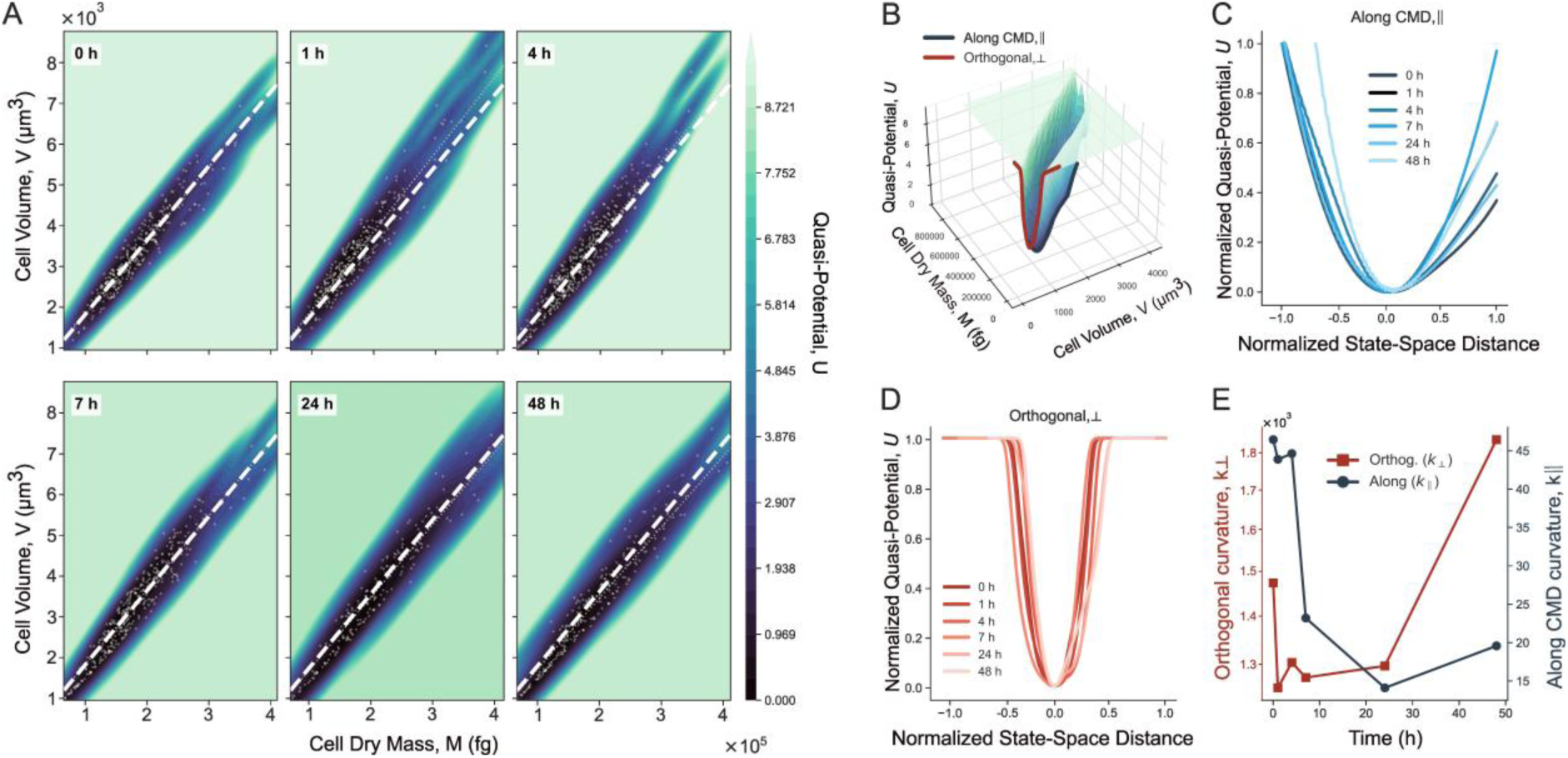
Quasi-potential landscapes reveal an anisotropic attractor underlying cell mass density homeostasis. (A) Empirical *U* reconstructed in mass–volume space during 48 h of recovery following osmotic shock. Dashed white lines indicate the target cell mass density (CMD; ρ₀ = 182.1 fg μm⁻³). (B) Three-dimensional representation of the baseline (0 h) landscape showing the principal axes orthogonal (red) and along (blue) the CMD manifold. (C,D) Normalized potential profiles extracted along the parallel (C) and orthogonal (D) directions over time (0–48 h). The potential remains broad along the parallel axis but sharply confined along the orthogonal axis throughout recovery. (E) Temporal evolution of the local dimensionless curvature (κ) at the quasi-potential minimum, calculated in normalized mass–volume state space. Orthogonal curvature (κ⊥) remains substantially greater than parallel curvature (κ∥) throughout recovery, indicating persistent anisotropic geometric confinement of cell states around the target density.

To determine whether this structural organization persists during the stress response, we reconstructed the quasi-potential landscapes (*U*) across the full 48-hour recovery period (**Fig. 2A**). Despite the pronounced population shifts induced by the osmotic shock, the fundamental topology of the landscape remained stable. The low-potential valley aligned with the CMD manifold persisted as an enduring geometric feature, demonstrating that density homeostasis is governed by a structure that is persistent over the measured recovery period rather than a transient physiological adaptation.

This attractor exhibits pronounced geometric anisotropy, defining two distinct geometric confinement regimes. By extracting the local potential profiles parallel and orthogonal to the CMD manifold, we mapped the normalized and dimensionless geometric confinement associated with state-space displacements (**Fig. 2C, 2D**). Along the CMD axis, the potential defines a wide, shallow basin, imposing minimal restrictions on coordinated mass-volume scaling. In sharp contrast, orthogonal displacements that alter cell mass density are opposed by steep potential walls, reflecting stronger restoring tendency. This strict directional asymmetry was preserved across all timepoints, maintaining tight density confinement even as the population relaxed from the initial perturbation.

To formally quantify this differential confinement, we calculated the local landscape curvature (κ) at the potential minimum for both principal axes (**Fig. 2E**). Throughout the entire recovery process, the orthogonal curvature (κ⊥=1.25–1.84×10^3^) remained consistently greater than the parallel curvature (κ∥ =14.1–46.4), corresponding to a ∼29–94-fold difference (Supplementary Table S1). Here, κ is dimensionless because both the quasi-potential and the relative state-space distance used for the curvature calculation are normalized, and therefore represents a geometric measure of attractor confinement rather than a physical mechanical stiffness. This pronounced geometric anisotropy mathematically defines the homeostatic process: the attractor architecture permits broad physiological variations in overall cell size and mass along the valley floor, while strongly constraining density-altering deviations. This inherent state-space geometry is consistent with the directed restoring dynamics that channel perturbed populations back toward the homeostatic setpoint.

### Optimal transport reveals restoring dynamics toward the density attractor

The anisotropic attractor identified in **Figure 2** suggests that recovery dynamics should be highly structured rather than random. Specifically, if the CMD manifold represents a low-potential valley within mass–volume state space, cellular trajectories should preferentially evolve along this manifold while minimizing excursions orthogonal to it. We therefore next sought to determine how populations move through the attractor landscape during recovery.

To reconstruct recovery dynamics, we applied optimal transport (OT) analysis to consecutive population distributions across the recovery time course. OT provides an estimate of the minimum-cost coupling of cellular states between successive timepoints and thereby enables inference of population-level flow within mass–volume space. The resulting OT-inferred transitions revealed a highly organized progression through state space that remained closely aligned with the CMD manifold throughout recovery (**Fig. 3A**). Despite substantial physiological changes following osmotic perturbation, population centroids evolved predominantly along the density-preserving valley identified in **Figure 2** rather than moving freely across the surrounding landscape. These observations suggest that recovery is strongly constrained by the underlying attractor geometry. Additionally, this approach reconstructs a continuous attractor displacement field, mapping the direction of population flow within the mass-volume state space (**Fig. 3B**).

**Figure 3.**
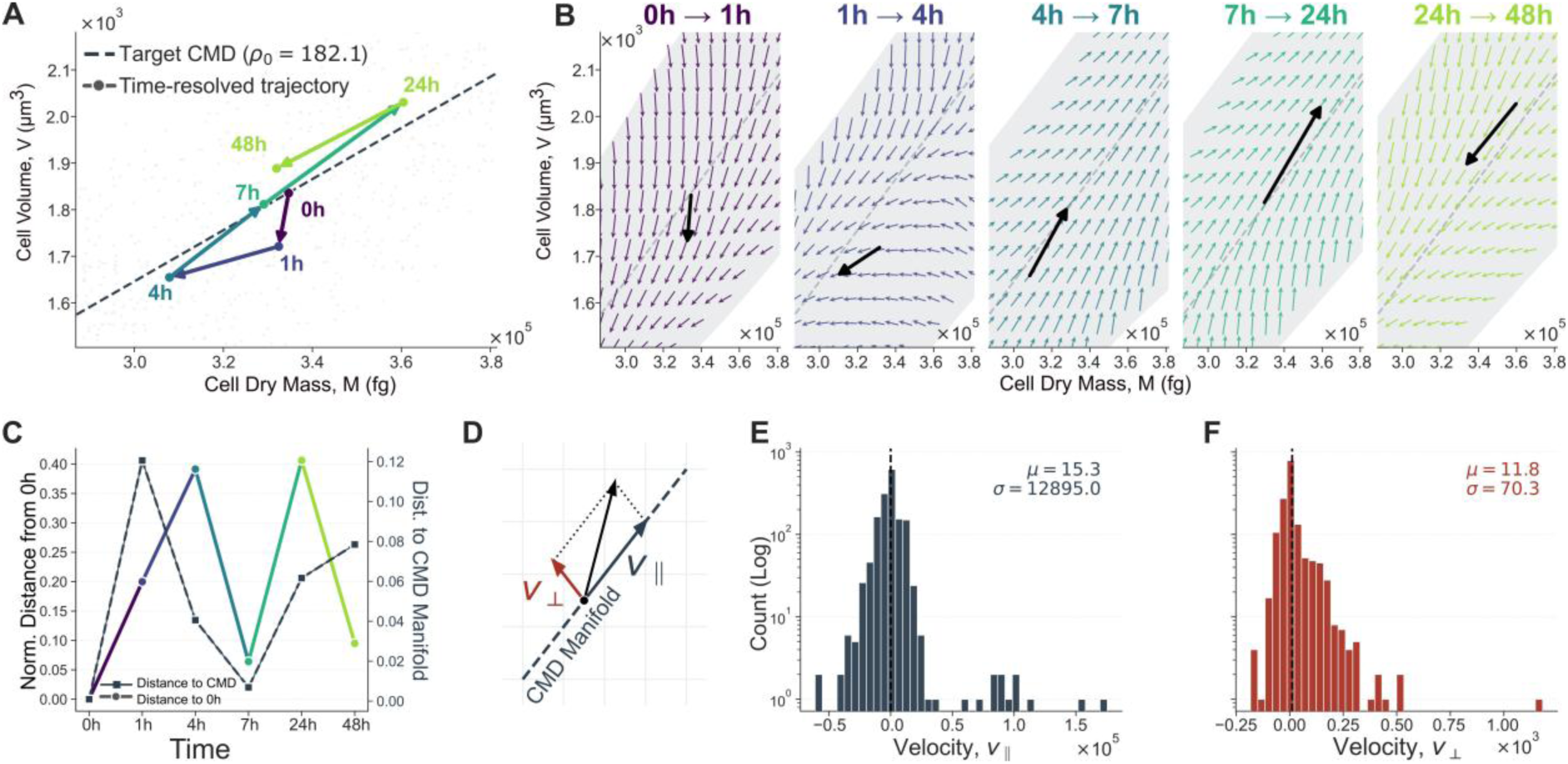
OT reveals anisotropic recovery constrained by the density attractor. (A) Population centroid trajectory in mass–volume space (0–48 h) relative to the target CMD manifold (dashed line, ρ₀ = 182.1 fg/μm³). (B) Time-resolved OT velocity vector fields mapping transition dynamics between consecutive timepoints. (C) Normalized Euclidean distance from the initial state (0 h) and orthogonal distance to the CMD manifold over time. (D) Schematic decomposition of velocity vectors into parallel (v∥) and orthogonal (v⊥) components relative to the CMD manifold. (E, F) Log-scaled distributions of the raw parallel and orthogonal OT-derived displacement rates. Because the two components retain the differing physical scales of mass and volume, their numerical dispersions are not directly compared here; the scale-normalized comparison is presented in Supplementary Fig. S2. Following normalization, displacement-rate variability is 1.79-fold greater along the CMD direction than orthogonal to it.

The resulting OT vector fields reveal a strongly structured, two-phase recovery pattern. The acute osmotic shock (0 to 1 h) drives a rapid, nearly orthogonal displacement across the state space, characterized by a massive volume contraction (ΔV_mean_ ≈ −830 μm³) with minimal dry mass loss (ΔM_mean_ ≈ −1.3×10⁴ fg), moving the population into a higher quasi-potential region. However, during the subsequent biological relaxation phase (1 to 48 h), the displacement vectors abruptly reorient. Rather than traversing the state space via direct Euclidean paths, the population flux is preferentially channeled along the low-potential valley. Population centroids evolve predominantly along the density-preserving manifold, maintaining a minimal orthogonal distance to the CMD manifold even as the Euclidean distance to the pre-shock state significantly varies (**Fig. 3C).**

To rigorously quantify these kinematic constraints, we decomposed the OT-derived velocity vectors, representing the state-space transition rates derived from the minimum-cost coupling between successive probability distributions, into components parallel (v∥) and orthogonal (v⊥) to the CMD manifold (**Fig. 3D**). This basis separates coordinated mass–volume fluctuations that preserve density from density-changing deviations that push cells away from the attractor valley. The raw projected displacement-rate distributions are shown in **Fig. 3E,F** (μ∥ = 15.3, σ∥ = 12,895.0; μ⊥ = 11.8, σ⊥ = 70.3). Because these projections retain the different physical scales of mass and volume, their numerical dispersions were not compared directly. After scaling mass and volume by their global standard deviations, the dispersion of OT- inferred displacement rates remained greater along the CMD direction than orthogonal to it (**Fig. S2**), with the standard deviation of the parallel component being 1.79-fold higher than that of the orthogonal component.

These findings provide the dynamical counterpart to the geometric anisotropy identified in **Figure 2**. The attractor not only shapes the static distribution of cellular states but constrains the inferred population flow during recovery. Together, the quasi-potential and optimal transport analyses indicate that density homeostasis is governed by a structured dynamical landscape: it permits substantial physiological flexibility along the CMD manifold while strongly restricting excursions that alter cell mass density.

### Coupled mass and volume feedback generate density-restoring dynamics

The attractor-constrained dynamics identified in **Figures 2** and **3** suggest the existence of a regulatory mechanism that senses density deviations and generates corrective responses. We therefore next asked whether density itself influences the physiological processes governing mass accumulation and volume regulation, thereby contributing to the restoring dynamics observed during recovery.

The restoring relationship between density deviation and subsequent density change suggests that recovery may be described by an effective feedback process. To quantify this relationship, we examined the interval-averaged rate of density change as a function of the initial density deviation for each OT-inferred state transition during the biological recovery phase (1–48 h), excluding the externally imposed 0–1 h osmotic perturbation (**Fig. 4A)**.

**Figure 4.**
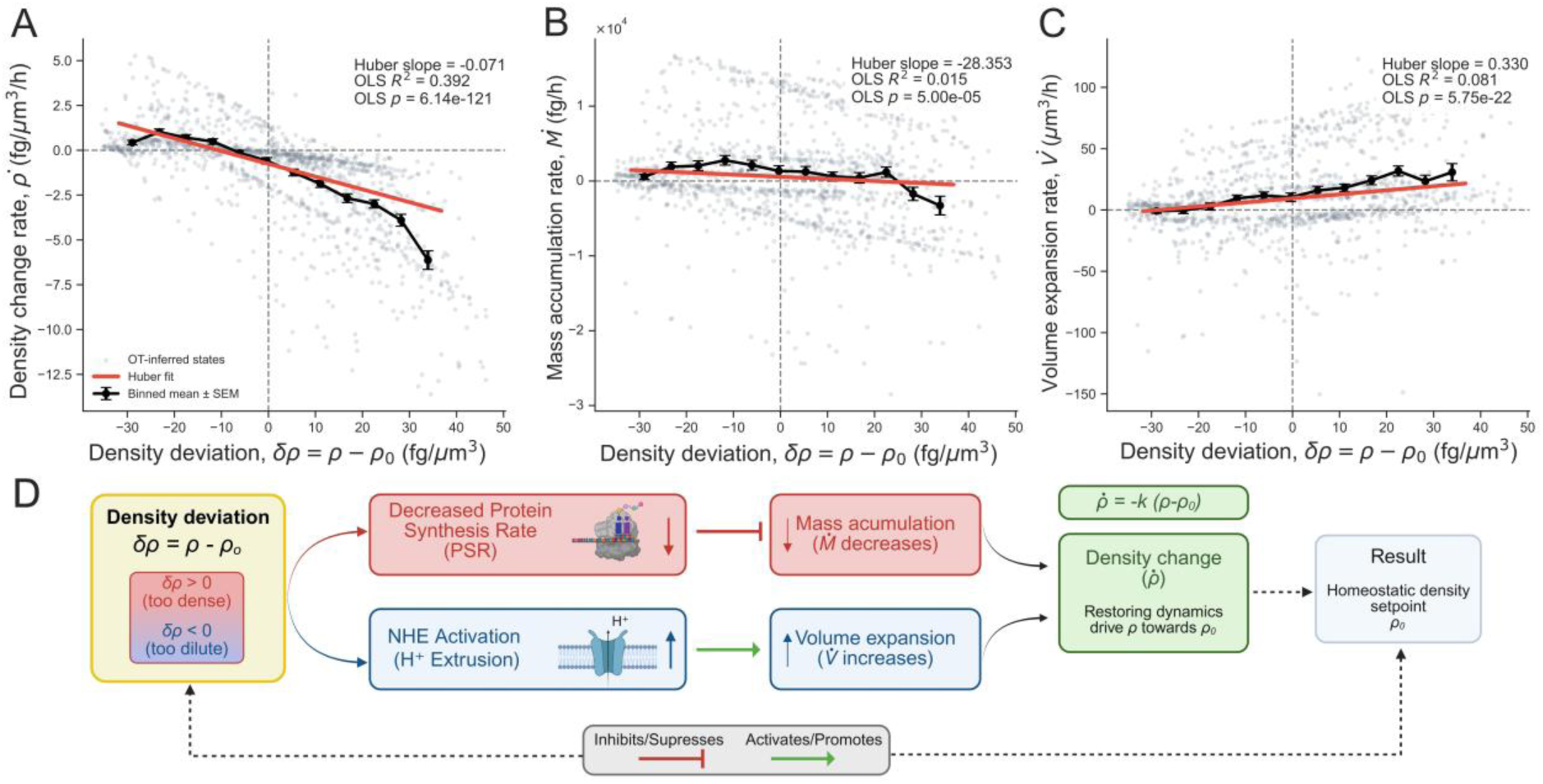
Coupled mass and volume dynamics are associated with density-restoring behavior. (A) Interval-averaged density change rate (ρ̇) as a function of density deviation from the homeostatic target (δρ = ρ - ρ₀) during the biological recovery phase (1–48 h); the acute 0–1 h osmotic perturbation was excluded. Gray points represent OT-inferred state transitions and black markers indicate binned means ± SEM. The red line shows the Huber regression (slope = −0.071); OLS statistics over the same analysis domain were R² = 0.392 and p = 6.14 × 10⁻¹²¹. (B,C) Relationships between density deviation and (B) dry-mass accumulation rate (Ṁ) and (C) volume expansion rate (V̇). Elevated density was weakly associated with reduced dry-mass accumulation (Huber slope = −28.353; OLS R² = 0.015, p = 5.00 × 10⁻⁵) and more consistently associated with increased volume expansion (Huber slope = 0.330; OLS R² = 0.081, p = 5.75 × 10⁻²²). (D) Proposed mechanistic interpretation of cellular density homeostasis informed by prior molecular studies.^14^ Density-dependent modulation of protein synthesis and NHE-mediated proton extrusion are shown as candidate mechanisms linking mass accumulation and volume regulation to the observed restoring dynamics.

This revealed a highly significant negative relationship (Huber slope = −0.071; OLS R² = 0.392, p = 6.14 × 10⁻¹²¹). Cells denser than the homeostatic target (δρ > 0) were associated with subsequent decreases in density, whereas cells below the target density (δρ < 0) were associated with subsequent increases. The negative restoring relationship was preserved across all individual recovery intervals, indicating that density deviations from the target density are systematically associated with corrective dynamics directed toward the homeostatic state. These observations provide quantitative evidence consistent with an effective density-restoring control law operating within the system.

To determine how mass and volume dynamics contribute to this restoring behavior, we separately examined the relationships between density deviation and dry-mass accumulation rate (Ṁ) and volume expansion rate (V̇). Elevated density showed an overall association with reduced dry-mass accumulation (Huber slope = −28.353; OLS R² = 0.015, p = 5.00 × 10⁻⁵; **Fig. 4B**), although this relationship varied across individual recovery intervals. In contrast, volume expansion exhibited a consistent positive association with density deviation (Huber slope = 0.330; OLS R² = 0.081, p = 5.75 × 10⁻²²; **Fig. 4C**), with positive slopes observed across all recovery intervals. Thus, hyper-dense states were associated with increased volume expansion and, more weakly, with reduced dry-mass accumulation, whereas lower-density states exhibited the opposite overall tendency. This error-dependent divergence is also evident in the time-resolved OT-inferred state transitions, which reveal opposed but coordinated kinematic responses across successive recovery intervals (Supplementary **Fig. S3**).

Taken together, these results indicate that density restoration is associated with coordinated changes in both mass and volume dynamics, with a particularly consistent density-dependent modulation of volume expansion and a weaker, temporally heterogeneous association with dry-mass accumulation. These relationships provide a physiological basis for the density-restoring behavior observed in **Figure 4A** and are consistent with the attractor-constrained recovery dynamics identified in **Figure 3**.

These coupled interactions suggest a specific, bipartite control architecture for cellular density homeostasis (**Fig. 4D**). Informed by prior mechanistic studies,^14^ we propose a framework where density deviations are associated with the physical or biochemical modulation of key metabolic pathways, decreasing protein synthesis rates (PSR) to halt mass accumulation, while activating membrane ion exchangers (e.g., NHE-mediated H⁺ extrusion) to drive osmotic volume expansion. Within this interpretation, robust density homeostasis (ρ̇ = −kδρ) emerges not from the independent tracking of mass or volume, but from their continuous, coordinated feedback regulation. This framework links the state-space geometry revealed in **Figure 2** with the recovery dynamics observed in **Figure 3** and provides a mechanistic basis for understanding how robust density homeostasis is achieved following perturbation.

### Density homeostasis suppresses fluctuations and encodes predictive information about future corrective behavior

The feedback architecture inferred in Figure 4 predicts that density homeostasis should actively suppress fluctuations and generate corrective responses whose magnitude depends on the underlying density error. We therefore next sought to quantify the performance of the inferred controller and determine whether density deviations contain predictive information regarding future cellular behavior.

As an initial measure of controller performance, we examined the evolution of density variance throughout recovery (**Fig. 5, left panel**). Density variance increased following osmotic perturbation, reaching a maximum at 4 h, and subsequently declined during recovery. Although variance remained above the pre-perturbation baseline at 48 h, the post-peak decrease was consistent with progressive suppression of density fluctuations around the empirical attractor.

**Figure 5.**
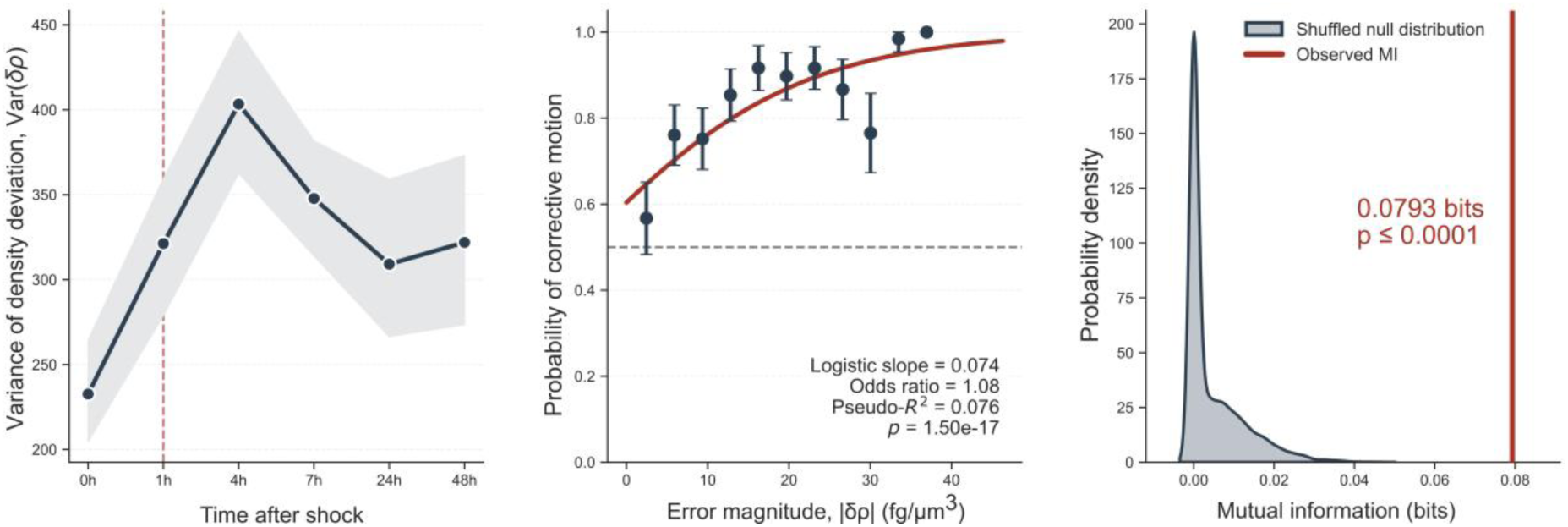
Density homeostasis suppresses fluctuations and encodes predictive information. (A) Temporal evolution of population density variance, Var(δρ), during the 48-h recovery. Density variance increased after osmotic perturbation, peaked at 4 h, and subsequently declined, while remaining above the pre-perturbation baseline at 48 h. Shaded regions denote bootstrap 95% confidence intervals (300 resamples). (B) Probability of directionally corrective motion as a function of absolute density error (|δρ|) during the biological recovery phase (1–48 h); the acute 0–1 h osmotic perturbation was excluded. Binned probabilities are shown with 95% binomial confidence intervals and the red curve indicates the logistic regression fit (slope = 0.074, odds ratio = 1.08, pseudo-R² = 0.076, p = 1.50 × 10⁻¹⁷). The dashed horizontal line indicates a probability reference of 0.5. (C) Mutual information (MI) between |δρ| and the subsequent binary corrective outcome. The observed MI was 0.0793 bits and exceeded the shuffled null distribution generated from 10,000 permutations (p ≤ 1.0 × 10⁻⁴), indicating predictive information about subsequent corrective behavior.

A hallmark of feedback regulation is that corrective action increases with the magnitude of the deviation. To test this in our system, we mapped the probability of a subsequent directionally corrective step against the absolute density error (|δρ|) during the biological recovery phase (**Fig. 5, middle panel**). Logistic regression revealed a highly significant monotonic relationship (slope = 0.074, odds ratio = 1.08, pseudo-R² = 0.076, p = 1.50 × 10⁻¹⁷), indicating that the odds of a corrective response increase with error magnitude. This positive relationship was also observed within each individual recovery interval. Thus, larger density deviations were increasingly associated with corrective responses directed toward the homeostatic setpoint, thereby reinforcing the restoring velocity field mapped in **Figure 3**.

Beyond corrective behavior itself, we asked whether the magnitude of the current density deviation contains information about subsequent regulatory behavior. Mutual information between |δρ| and the binary corrective outcome was 0.0793 bits (**Fig. 5, right panel**), exceeding the shuffled null distribution (10,000 permutations; p ≤ 1.0 × 10⁻⁴). These results indicate that the current density-error magnitude contains predictive information about the directionality of the subsequent OT-inferred density response.

Together, these findings demonstrate that cellular density homeostasis exhibits several hallmarks of an effective dynamical control system. The inferred controller suppresses fluctuations, increases corrective effort as deviations grow larger, and encodes significant predictive information regarding future responses. These properties provide a quantitative framework for understanding how populations maintain robust density regulation despite pervasive cellular heterogeneity and environmental perturbation, completing the progression from attractor geometry (**Fig. 2)** and recovery dynamics (**Fig. 3**) to control architecture (**Fig. 4**) and controller performance (**Fig. 5**).

## Discussion

In this study, we revisited single-cell mass–volume measurements acquired during recovery from osmotic perturbation within the framework of dynamical systems theory, optimal transport, and information theory. This perspective revealed several previously unrecognized features of cellular density homeostasis. Specifically, we demonstrate that density regulation is associated with a persistent anisotropic attractor embedded in mass–volume state space, that recovery follows highly structured trajectories constrained by the geometry of this attractor, that density deviations are coupled to coordinated feedback acting on both mass accumulation and volume regulation, and that density errors contain significant predictive information about future corrective behavior. Together, these findings suggest that cellular density homeostasis can be understood as an emergent dynamical control process rather than simply as a consequence of coupled changes in mass and volume.

Our previous work established that mammalian cells actively restore density following osmotic perturbation through coordinated regulation of dry mass and volume, identifying key molecular mechanisms and proposing a macroscopic proportional-integral-derivative (PID) control model. However, the large-scale kinetic analyses in that study primarily relied on calculating density as the ratio of population-averaged mass to population-averaged volume, derived from separate quantitative phase microscopy and fluorescence eXclusion measurements. In the present study, optical diffraction tomography enables simultaneous single-cell estimates of dry mass and volume, allowing us to resolve their joint organization and dynamics in mass–volume state space. The reference CMD obtained here differs modestly from that reported previously, reflecting differences in the analysis pipeline^30^ (see Methods).

By reanalyzing the system from a state-space perspective, we resolve single-cell dynamical trends and transition trajectories that were not accessible through previous population-level analyses. This approach demonstrates that density regulation is organized around a low-dimensional attractor that constrains both the geometry of cellular states and the structured population-level flow connecting successive distributions, providing a new layer of biological insight into how density errors are associated with subsequent corrective behavior.

A central finding is that density regulation is governed by an anisotropic attractor, challenging the classical notion of homeostasis as a convergence toward a rigid setpoint.^31^ Instead, cells occupy an extended low-potential valley corresponding to a preferred target density. This design is biologically imperative: because cells continuously grow,^32^ remodel their proteomes,^33^ alter water content,^34^ and progress through the cell cycle, strict regulation of either mass or volume alone would be incompatible with normal physiology.^17^ The anisotropic geometry naturally solves this by strongly restricting density-altering deviations while permitting broad density-preserving fluctuations. The markedly greater curvature orthogonal to the valley than along it (approximately 29–94-fold across recovery) provides a quantitative measure of this anisotropy. This effectively balances robust homeostatic control with the physiological flexibility required for growth and remodeling.

The optimal transport analyses^27^ further revealed that recovery dynamics are strongly shaped by this attractor geometry. Rather than exhibiting unconstrained motion through mass–volume space, population trajectories remained closely aligned with the CMD manifold throughout recovery. This observation suggests that the attractor is not merely a statistical feature of the observed distributions but reflects an underlying dynamical structure that channels cellular behavior. Although optimal transport does not reconstruct exact single-cell trajectories, the inferred population-level flows provide a compelling picture of how homeostatic recovery unfolds within the constraints imposed by the state-space geometry. Together, the geometric and dynamical analyses support a view of density homeostasis as a structured navigation problem occurring on a low-dimensional manifold embedded within a higher-dimensional physiological landscape.

A second major finding, consistent with our previous study, is that density restoration appears to emerge from coordinated feedback acting on both mass accumulation and volume regulation. Here, density deviations were consistently associated with volume expansion and showed a weaker, interval-dependent association with mass accumulation. Although the present state-space analysis does not directly identify the underlying molecular circuitry, these relationships are consistent with mechanisms identified in our previous work, in which elevated mass density promoted volume recovery through activation of membrane ion transport, including the Na+/H+ exchanger (NHE), while suppressing mass accumulation through density-dependent changes in nucleocytoplasmic transport and protein synthesis. These mechanisms are also consistent with broader observations linking macromolecular crowding to reduced translational efficiency^35^ and osmotic adaptation to ion transport and cell-volume regulation.^36,37^ From this perspective, density homeostasis may not require a single dedicated density-sensing pathway, but may instead emerge from the coordinated action of multiple physiological processes that collectively generate an effective density-restoring controller. ^38^

This interpretation highlights an important conceptual distinction between mechanism and control. The control law inferred here should not be viewed as a molecular pathway but rather as a coarse-grained description of how the system responds to perturbation. Similar effective control laws have proven useful in fields ranging from neuroscience^11,39^ to ecology, where complex networks of interacting components give rise to relatively simple emergent behaviors at larger scales. The density-restoring law identified in this work therefore provides a quantitative description of homeostatic regulation without requiring prior knowledge of the underlying molecular circuitry.

Perhaps the most intriguing result emerged from the information-theoretic analysis. Homeostasis is often evaluated in terms of fluctuation suppression or restoration of average behavior. However, suppression alone does not necessarily imply active regulation.^40^ The observation that density-error magnitude contains 0.0793 bits of statistically significant predictive information about subsequent corrective behavior indicates a measurable dependence between the current state and the direction of the inferred response. Rather than establishing density as a causal control signal, this result supports its interpretation as an informative state variable associated with subsequent corrective action. This finding is consistent with broader perspectives that view biological regulation as an information-processing problem, in which the current physiological state can carry information relevant to subsequent regulatory responses. ^41,42^

More broadly, the analytical framework developed here may prove applicable well beyond density regulation. The combination of state-space reconstruction, optimal transport, and information-theoretic analysis provides a general strategy for inferring attractor geometry, restoring dynamics, and feedback structure directly from single-cell measurements. While we focused on osmotic recovery and cellular density homeostasis, similar approaches could be applied to diverse biological systems including cell-size control,^43^ metabolic adaptation,^8^ differentiation dynamics,^27^ immune activation,^44^ microbial stress responses,^45^ and tissue-level^46^ homeostatic processes. In this sense, the present study contributes not only to our understanding of density regulation but also to the broader challenge of reconstructing biological control systems from high-dimensional experimental data.

Several limitations should be acknowledged. First, the inferred control laws represent effective population-level descriptions and do not directly identify the molecular pathways responsible for density restoration.^47^ Second, OT infers a minimum-cost population-level coupling and therefore provides an approximation of the underlying dynamics rather than exact single-cell trajectories. Third, the present study focuses on a single perturbation paradigm and cellular system. Future work will be required to determine the extent to which the inferred attractor geometry and control architecture generalize across cell types, physiological conditions, and forms of environmental stress.

In conclusion, our results suggest that cellular density homeostasis can be understood through the unifying framework of dynamical systems, control theory, and information processing. By revealing how attractor geometry, restoring dynamics, and coupled feedback mechanisms interact to stabilize cell mass density, this work provides a quantitative description of homeostatic regulation as an emergent control process. More broadly, these findings suggest that homeostatic variables may be understood not solely through their absolute values, but through the attractor geometries, restoring dynamics, and feedback architectures that stabilize them. We anticipate that similar approaches will prove valuable for uncovering the organizing principles governing biological homeostasis across scales.

## Materials and Methods

### Cell culture, osmotic perturbation, and quantitative phase imaging

The single-cell mass and volume measurements analyzed in this study were obtained from a previously published quantitative phase imaging (QPI) dataset investigating cellular recovery following osmotic perturbation. Experimental procedures, cell culture conditions, osmotic treatment protocols, image acquisition parameters, and extraction of single-cell dry mass and volume measurements have been described previously.^14^

Briefly, NIH3T3 cells were subjected to acute osmotic perturbation and imaged at multiple recovery timepoints (0, 1, 4, 7, 24, and 48 h) using commercial Optical Diffraction Tomography instrument (HT-2H, TomoCube Inc., Korea). These measurements formed the basis for all subsequent analyses.

### Cell mass density and density deviations

Cell mass density (CMD) was defined as

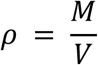

where M denotes cellular dry mass and V denotes cellular volume. M and V were calculated directly from 3D refractive-index tomograms (with voxel size of 0.095 × 0.095 × 0.190 μm³). 2D masks were created using TomoAnalysis (TomoCube, Korea) software and manually proof-read and reviewed against the corresponding cell images before the same downstream 3D segmentation procedure (CellSNAP) was applied, as applied in previous studies^30^ Briefly, CellSNAP utilizes 2D masks to spatially gate the 3D tomograms, followed by segmentation using a threshold of 1.36 and subsequent connected-component-based noise removal.) Dry mass was calculated by integrating the RI difference between the cell and the background media (RI= 1.3364) utilizing a specific RI increment (α) of 0.18 ml/g, using the following formula from literature^48,49^

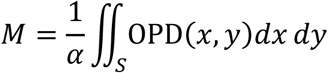

Where OPD is calculated as

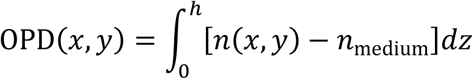

Prior to analysis, cells with volumes exceeding 6000 μm³ or dry masses exceeding 1,000,000 fg were excluded as outliers.

The target density (ρ₀) was estimated from the baseline cells (t = 0 h) as the slope of a robust Huber regression of dry mass against volume, constrained to pass through the origin.

Density deviations were calculated as

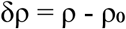

such that positive values indicate cells denser than the target state and negative values indicate cells with lower-than-target density. This manifold serves as the density-preserving axis within mass–volume state space.

The target density obtained here differs modestly from that reported in our previous work^14^, reflecting differences in dry-mass calculation pipelines used for the respective analyses. Importantly, the same processing pipeline and internally estimated reference CMD were applied consistently across all timepoints and downstream analyses in the present study.

### Empirical quasi-potential landscape reconstruction

To characterize the geometry of density homeostasis, empirical quasi-potential landscapes were reconstructed directly from the observed population distributions in mass–volume space. For each recovery timepoint, the joint probability density P(M,V) was estimated using two-dimensional kernel density estimation (KDE).

The quasi-potential landscape was then computed as

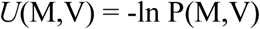

where low values of *U* correspond to highly populated states and high values correspond to sparsely occupied states.

Landscape surfaces were evaluated on a regular grid spanning the observed mass–volume range. Specifically, the probability density was estimated using a Gaussian kernel density estimation (KDE) implemented via SciPy (scipy.stats.gaussian_kde). Bandwidth was selected based on Scott’s rule, scaled by an empirical factor of 1.2 to slightly smooth individual cell noise. Landscape surfaces were evaluated on a regular 150 × 150 grid spanning the global physical limits of the observed mass and volume. To compute the quasi-potential, a small constant (10⁻¹²) was added to prevent logarithmic undefined values, and the density was normalized to its local maximum, ensuring the basin minimum sits at U = 0. For robust 2D visualization, quasi-potential values were clipped at the global 99th percentile, while 1D potential profiles (slices) were Min-Max normalized between 0 and 1 for visualization.

### Quantification of attractor geometry and anisotropy

To quantify local attractor geometry, quasi-potential profiles (1D slices) were extracted along two orthogonal directions. Rather than imposing a fixed geometric axis, these directions were determined dynamically for each timepoint using Principal Component Analysis (PCA) on mass and volume coordinates independently normalized to their global observed ranges. The same global scaling was used across all timepoints. The first principal component defined the direction of maximal population variance, approximately aligned with the CMD manifold, while the second principal component defined the corresponding orthogonal direction. The calculated parallel and orthogonal vectors were subsequently mapped back onto the physical mass–volume space to extract the corresponding quasi-potential profiles (Figure S1).

Potential slices were centered on the minimum of the quasi-potential valley and sampled symmetrically in both directions. For visualization, the 1D potential profiles shown in Figure 2C,D were Min-Max normalized between 0 and 1.

To estimate the local curvature (κ), we applied a dynamic, symmetric fitting window comprising the central 30% of the extracted profile points (15% on each side of the quasi-potential minimum). Curvature was evaluated with respect to the dimensionless distance x in the normalized mass–volume state space. Within this restricted local basin, the quasi-potential data were modeled via an ordinary least-squares quadratic fit:

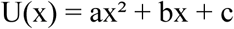

The local curvature was then analytically extracted as the second derivative of the fitted polynomial:

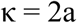

Because both *U* and x are dimensionless, the fitted coefficients and κ are also dimensionless. Here, κ represents a geometric measure of local quasi-potential confinement rather than a physical mechanical stiffness. Parallel (κ∥) and orthogonal (κ⊥) curvatures were tracked throughout recovery to quantify attractor anisotropy

### Optimal transport analysis

Population-level recovery dynamics were reconstructed using entropically regularized optimal transport (OT).^25,27^ For each pair of consecutive timepoints, empirical distributions of cells in mass–volume space were treated as discrete probability measures. By computing the minimum-cost coupling between successive distributions, OT enables the inference of population-level flow without assuming single-cell tracking. To ensure scale invariance and robust convergence, state-space variables were standardized by their standard deviations, and the cost matrix was defined using the median-normalized squared Euclidean distance. The OT plan was computed via a custom implementation of the Sinkhorn-Knopp algorithm^50^ (entropic regularization parameter λ = 0.07, maximum iterations = 1500, convergence tolerance = 10⁻⁹). Rather than employing state-space clustering, transition vectors were inferred directly for all individual observed states using the barycentric projection of the transport plan, yielding a dense, high-resolution vector field of expected mass-volume velocities. Population centroids were calculated as the mean coordinates of the population at each timepoint to visualize the macroscopic recovery trajectory.

### State-space flux field visualization

To visualize the continuous macroscopic flow of the population throughout recovery, the discrete, single-cell OT displacement rates were interpolated onto a uniform spatial grid using linear Delaunay interpolation (scipy.interpolate.griddata). To strictly prevent the extrapolation of flows into biologically unoccupied regions of the state space, the vector field was masked using a local probability density threshold. Specifically, a two-dimensional Gaussian kernel density estimate (KDE) was computed for the observed cellular coordinates, and interpolated flow vectors were only rendered in regions where the local density exceeded 5% of the maximum landscape density.

### Velocity decomposition

To quantify how recovery dynamics relate to attractor geometry, OT-derived displacement vectors were decomposed into components parallel and orthogonal to the CMD manifold. For each displacement vector (v),

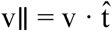

and

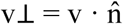

where t^ denotes the unit tangent vector of the CMD manifold and n^ denotes the corresponding orthogonal unit vector. Distributions of v∥ and v⊥ were compared to quantify the relative freedom of motion along density-preserving and density-changing directions. For direct comparison of the parallel and orthogonal displacement-rate distributions, mass and volume components were scaled by their respective global standard deviations, and the CMD-aligned tangent and orthogonal directions were defined in the same normalized state space before projection (Supplementary Fig. S2).

### Density-restoring law and feedback analysis

To identify effective feedback relationships, density deviation was related to subsequent interval-averaged changes in density, dry mass, and volume. The analysis was restricted to the biological recovery phase (1–48 h); the acute 0–1 h osmotic perturbation was excluded.

For each OT-inferred transition from a source state at time t to its barycentric destination at time t + Δt, the density change rate was defined as

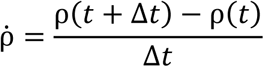

and similarly

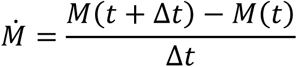

and

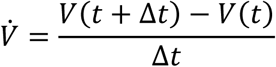

Density deviation was calculated relative to the common target density ρ₀ used throughout the study. Before regression analysis, extreme OT-inferred transitions were removed using a simultaneous 0.5th–99.5th percentile filter applied to δρ, ρ̇, Ṁ, and V̇; a transition was retained only when all four quantities fell within their respective percentile ranges.

To visualize state-dependent trends, OT-inferred source states were discretized into 12 equally spaced bins spanning the 3rd to the 97th percentile of the observed density deviation. Within each bin, the central tendency was estimated as the sample mean, with error bars representing the SEM. The same 3rd–97th percentile density-deviation domain was used for all regression analyses. Robust linear trends were estimated using Huber regression, implemented in scikit-learn, to reduce the influence of extreme observations.^51^ The slope reported for the robust fit corresponds to the Huber regression. For conventional inferential statistics, ordinary least-squares (OLS) regression was performed over the same observations and domain, and the associated coefficient of determination (R²) and two-sided p-value for the slope were reported.

### Controller performance analysis

Population-level density variance was calculated at each recovery timepoint as Var(δρ). Uncertainty was estimated using a non-parametric bootstrap with 300 resamples, and 95% confidence intervals were defined by the 2.5th and 97.5th percentiles of the bootstrap distribution.

Corrective behavior was analyzed only during the biological recovery phase (1–48 h); the acute 0–1 h osmotic perturbation was excluded. OT-inferred transitions were classified as directionally corrective when the interval-averaged density-change rate (ρ̇) opposed the density deviation (δρ), such that −δρ·ρ̇ > 0. This criterion indicates that the inferred density-change rate points toward the homeostatic setpoint, without requiring the finite transition to end closer to the setpoint in absolute distance.

Before analysis, extreme dynamic states were removed using a simultaneous 0.5th–99.5th percentile filter applied to δρ and ρ̇. The probability of a corrective transition was then modeled as a function of the absolute density deviation (|δρ|) using logistic regression implemented with the statsmodels^52^ Logit maximum-likelihood estimator and an explicit intercept term. The logistic slope, odds ratio, McFadden pseudo-R², and p-value for the slope were obtained directly from the fitted model. For visualization, corrective probabilities were calculated in 11 equally spaced bins spanning the 2nd to the 98th percentile of |δρ|; error bars represent 95% confidence intervals calculated using the standard binomial approximation.

### Information-theoretic analysis

To quantify predictive information contained within density deviations, MI was calculated between the absolute density-error magnitude and the discrete binary outcome of subsequent corrective behavior. Because this analysis involves mixed continuous and discrete variables, mutual information was estimated using the nearest-neighbor algorithm adapted for continuous-discrete data as described by Ross in 2014,^53^ implemented via the scikit-learn library. The nearest-neighbor parameter was strictly set to k = 5.

To determine the statistical significance of the observed mutual information, an empirical null distribution was constructed by generating 10,000 random permutations of the target variable, effectively shuffling corrective outcomes while preserving the marginal distributions of both variables. For each shuffle, MI was recomputed, and the observed MI value was compared against this empirical distribution to estimate statistical significance.

### Statistical analysis

All computational analyses, optimal transport modeling, and statistical testing were performed in Python (version 3.12.8). Data manipulation and numerical operations were conducted using the pandas (version 2.3.2) and NumPy (version 2.1.3) libraries. Statistical distributions, spatial algorithms, and ordinary least-squares regressions were computed using SciPy (version 1.16.2). Robust regression and mutual-information estimation were implemented using scikit-learn (version 1.7.2), whereas logistic regression was performed using statsmodels (version 0.14.6). Data visualization was generated using matplotlib (version 3.10.6) and seaborn (version 0.13.2). Results are presented as mean ± SEM unless otherwise indicated. For regression analyses, two-sided p-values were reported from ordinary least-squares or logistic regression, as specified. Mutual-information significance was assessed using permutation testing. Unless otherwise indicated, statistical significance was defined as p < 0.05.

## Acknowledgments

We thank Jinyu Fu and Sean X. Sun for the samples and collaboration. This work was supported by the National Institutes of Health (NIH) Maximizing Investigators’ Research Award (MIRA, R35GM149272 to I.B.) and the Air Force Office of Scientific Research Biophysics Program (FA9550-22-1-0334 to I.B.); V.A.R. acknowledges funding from the European Union’s Horizon Europe research under the Marie Skłodowska-Curie Actions (MSCA), RAMPART project (Grant Agreement No. 101209772).

## Supplementary Figures

**Figure S1.**
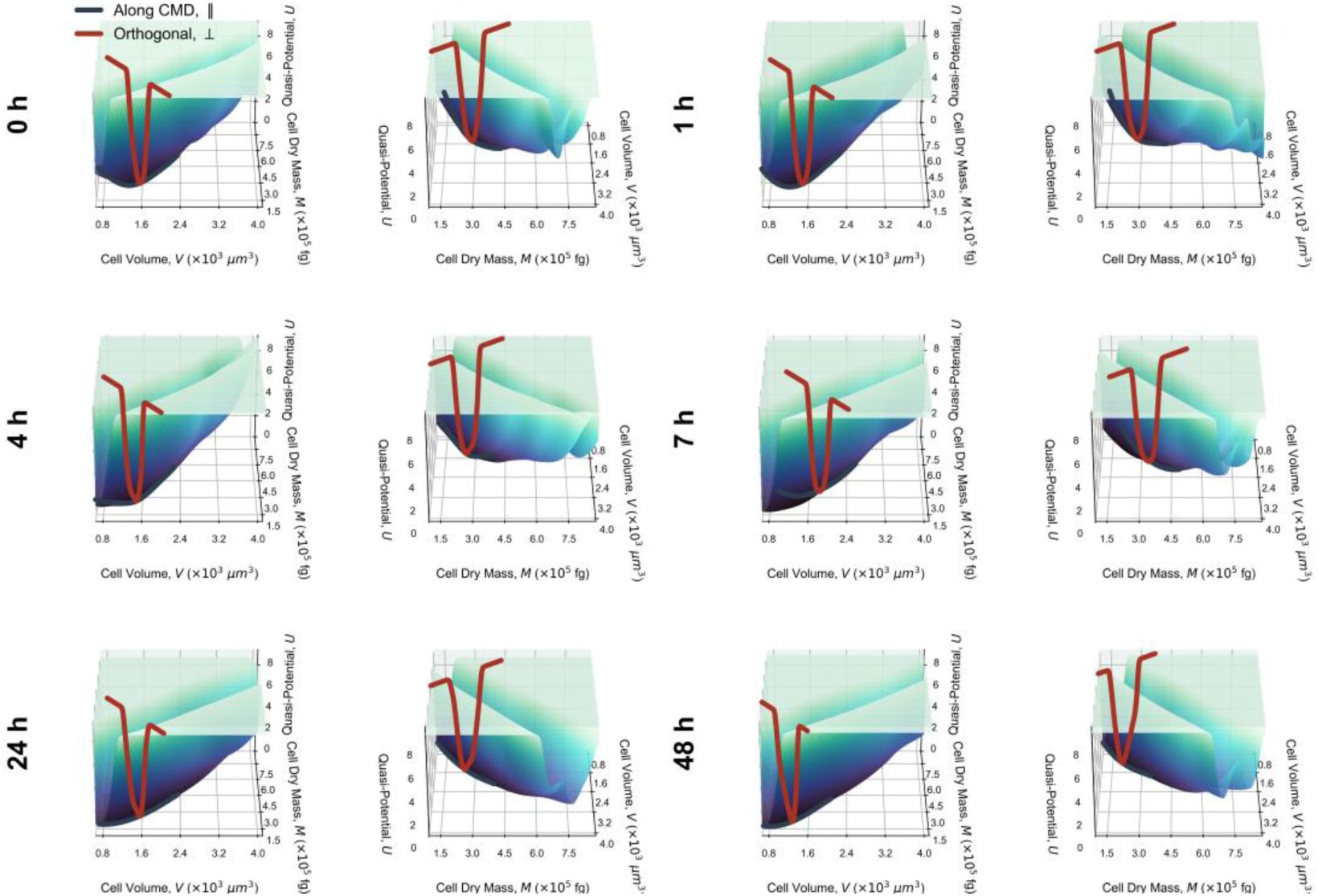
Three-dimensional geometric validation of quasi-potential landscapes using PCA. Topographic rendering of empirically reconstructed density attractors within the physical mass-volume state space throughout the recovery process (0 – 48 h). For each timepoint, quasi-potential surfaces (*U*) are displayed from two orthogonal perspectives (frontal and side views), based on PCA analysis. The overlaid trajectories visually confirm that the PCA-derived directions approximately aligned with the CMD manifold (Along CMD, dark blue) and orthogonal to it (Orthogonal, maroon) accurately bisect the basin of the homeostatic valley without spatial distortion.

**Supplementary Figure S2.**
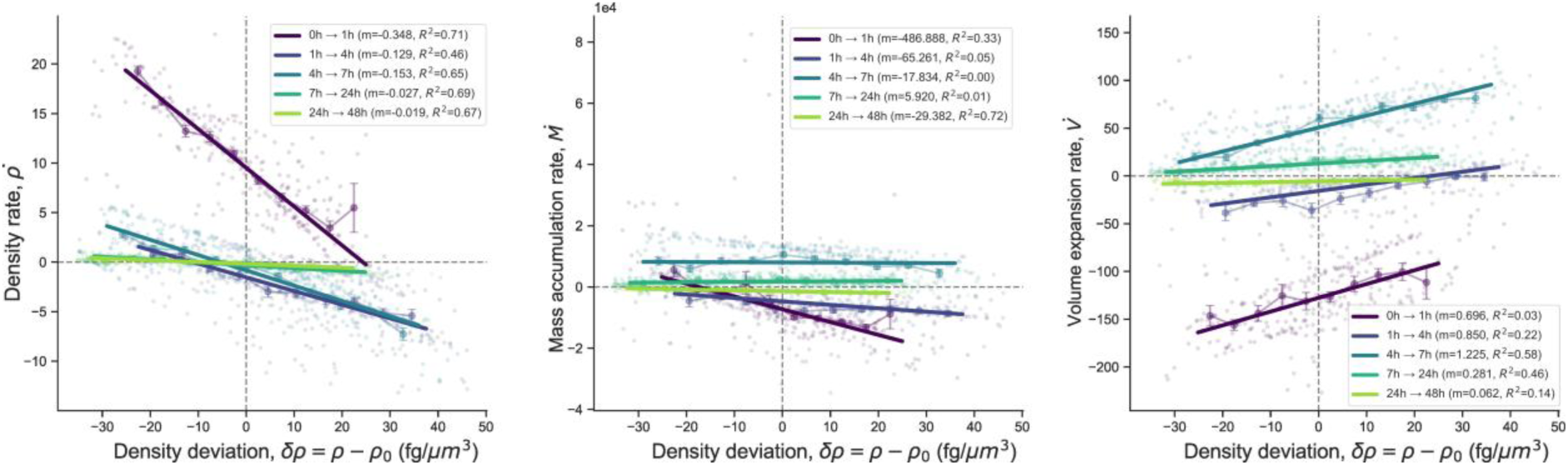
Normalized OT displacement-rate distributions along and orthogonal to the CMD manifold. (A,B) Distributions of OT-inferred displacement rates projected along the CMD direction (A, ϕ∥) and the orthogonal direction (B, ϕ⊥) after scaling cell volume and dry mass by their global standard deviations. Dashed lines indicate the mean values, and μ and σ denote the mean and standard deviation, respectively. The normalized parallel component exhibited a broader distribution than the orthogonal component (σ∥ = 0.139 h⁻¹ and σ⊥ = 0.078 h⁻¹), corresponding to a 1.79-fold higher standard deviation along the CMD direction.

**Figure S3.**
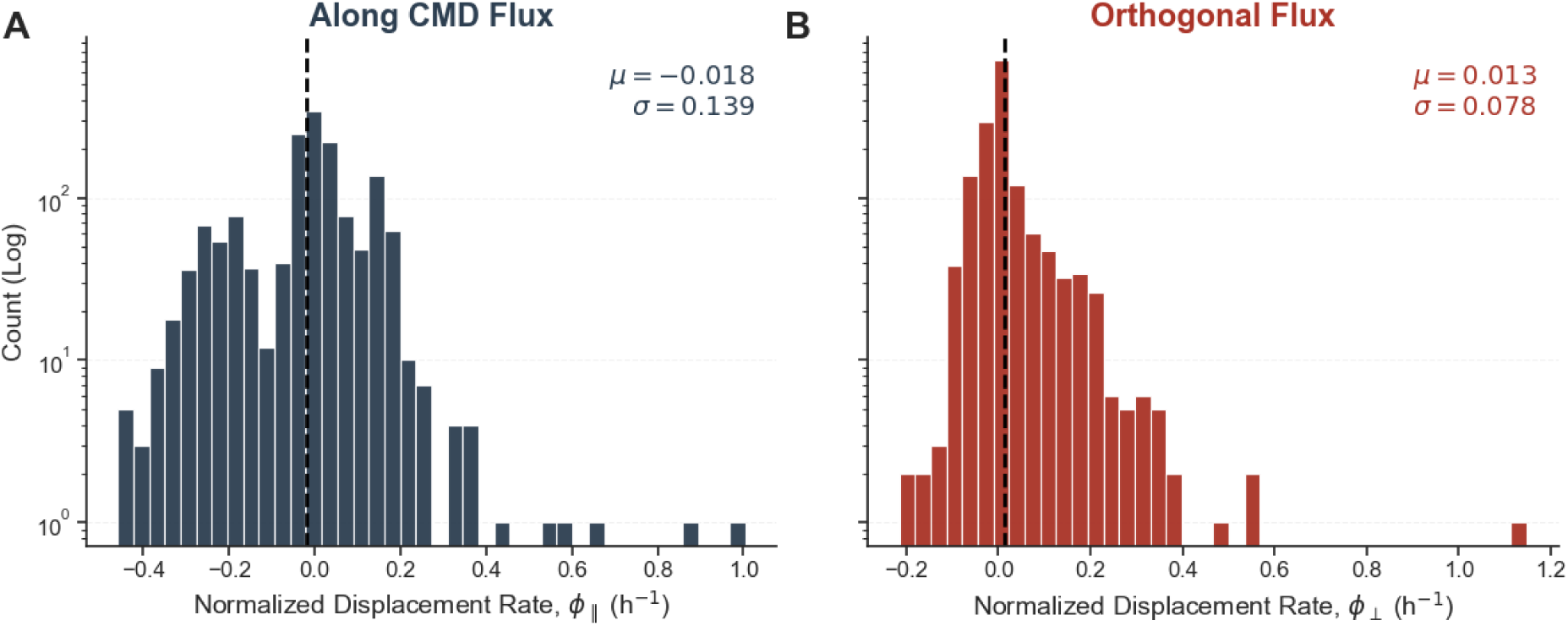
**Time-resolved analysis of density-dependent kinematic rates across the recovery period**. Scatter plots illustrating the relationship between initial density deviation (δρ = ρ - ρ₀) and the subsequent physiological rates for all measured optimal transport (OT) state transitions, stratified by temporal interval (0 h → 1 h, 1 h → 4 h, 4 h → 7 h, 7 h → 24 h, and 24 h → 48 h). (Left) Interval-averaged density change rate (ρ̇). (Middle) Mass accumulation rate (Ṁ). (Right) Volume expansion rate (V̇). Individual OT-derived state transitions are plotted as semi-transparent points, color-coded by their respective time interval. Binned means ± SEM are overlaid as opaque connected markers for each specific period. Solid lines represent independent linear regression fits for each time interval. The inset legends provide the extracted linear regression slope (m) and the coefficient of determination (R²) for each temporal transition.

**Supplementary Table S1.**
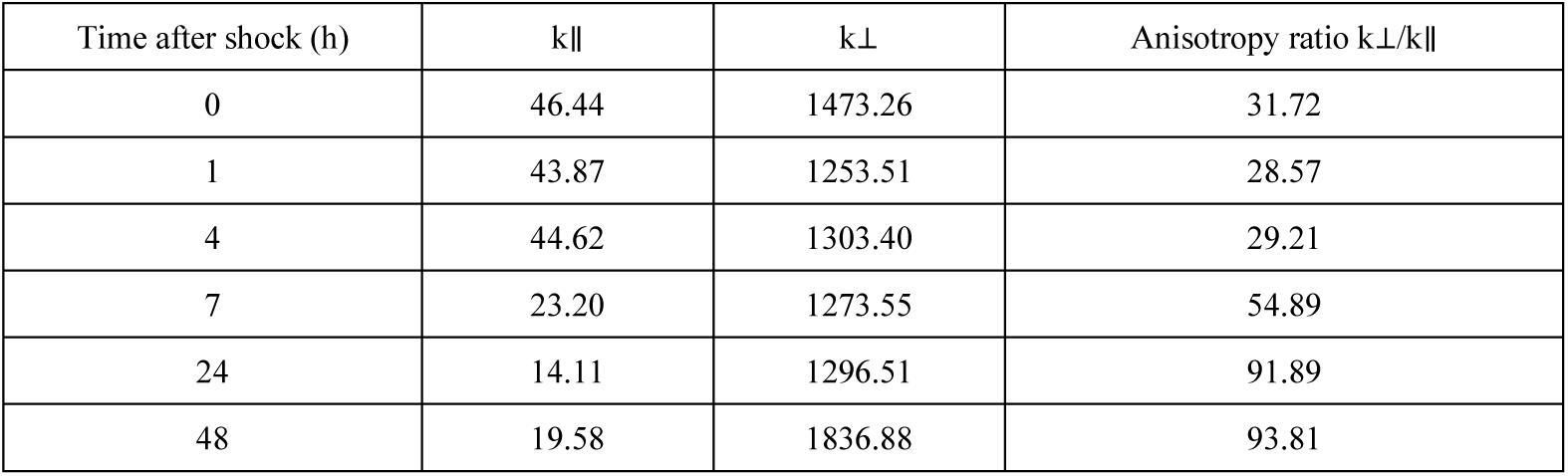
Local quasi-potential curvature and anisotropy during osmotic recovery. Local curvatures parallel (k∥) and orthogonal (k⊥) to the principal attractor directions were calculated from quadratic fits around the quasi-potential minimum. The anisotropy ratio was calculated as k⊥/k∥. The target cell mass density used throughout the analysis was ρ₀ = 182.1 fg μm⁻³.

